# Cholesterol Synthesis Pathway Genes in Prostate Cancer are consistently downregulated when tissue confounding is minimized

**DOI:** 10.1101/220400

**Authors:** Morten Beck Rye, Helena Bertilsson, Maria K. Andersen, Kjersti Rise, Tone F. Bathen, Finn Drabløs, May-Britt Tessem

## Abstract

The relationship between cholesterol and prostate cancer has been extensively studied for decades, where high levels of cellular cholesterol are generally associated with cancer progression and less favorable outcomes. However, the role of in vivo cellular cholesterol synthesis in this process is unclear, and data on the transcriptional activity of cholesterol synthesis pathway genes in tissue from prostate cancer patients are inconsistent. A common problem with cancer tissue data from patient cohorts is the presence of heterogeneous tissue which confounds molecular analysis of the samples. In this study we present a method to minimize systematic confounding from stroma tissue in seven patient cohorts consisting of 1713 prostate cancer and 230 normal tissue samples. When confounding was minimized, differential gene expression analysis over all cohorts showed robust and consistent downregulation of nearly all genes in the cholesterol synthesis pathway. Additional analysis also identified cholesterol synthesis as the most significantly altered metabolic pathway in prostate cancer. This surprising observation is important for our understanding of how prostate cancer cells regulate cholesterol levels in vivo. Moreover, we show that tissue heterogeneity explains the lack of consistency in previous expression analysis of cholesterol synthesis genes in prostate cancer.

## Introduction

Increased cholesterol levels in enlarged prostates and prostate cancer have been observed for decades^1-3^, and extensive research has suggested that cholesterol have a role in prostate cancer growth and progression^3-5^. Cholesterol homeostasis is important for cell viability, and is dynamically regulated by a balance between synthesis, uptake, efflux and storage of cholestero^l4,6-9^. For cellular cholesterol synthesis, the conversion of 3-hydroxy-3-methylglutaryl coenzyme A (HMG CoA) to mevalonate is the first rate limiting step, which is followed by over 20 flux controlling enzymatic reactions before cholesterol is synthesized as the final product. In prostate cancer cell-lines, elevated activity of the cholesterol synthesis pathway supports cancer growth and aggressiveness^10-16^. This has led to the general view that increased cholesterol synthesis in prostate cancer cells contributes to cellular accumulation of cholesterol and prostate cancer growth. A diet high in fat and cholesterol increase the risk of prostate cancer, while statins directly targeting the cholesterol synthesis pathway are associated with improved clinical outcome (reviewed in^17^). This is generally taken as support for the relevance of increased cholesterol synthesis *in vivo*. This notion was also in line with a recent study showing increased activity of the cholesterol synthesis enzyme squalene monooxygenase (*SQLE*) in lethal prostate cancer^18^. Accordingly, one would expect that genes in the cholesterol synthesis pathway are upregulated when prostate cancer is compared to normal tissue. However, transcriptional changes in cholesterol genes are rarely highlighted when such comparisons are performed in large patient cohorts.

We hypothesized that this is due to influence of confounding tissue components present in the samples. Gene expression analysis in human tissue is challenged by the highly heterogeneous tissue composition in each sample^19,20^. The standard way to account for such heterogeneity is to incorporate tissue type percentages from histopathology during the analysis. Although confounding due to tissue composition is generally acknowledged, data from histopathology are missing in most publicly available patient cohorts, which may bias the molecular analyses. In prostate cancer, the presence of stroma tissue is shown to hide underlying molecular features in a differential analysis^21,22^. Prostate tissues are usually histopathologically divided into benign epithelium, stroma tissue and prostate cancer. It is previously shown that the different number of tissue types present in prostate cancer (three tissue types) and in normal samples (two tissue types) leads to a systematic sampling bias of increased stroma content in the normal samples^23,24^. This confounds differential analysis when cancer and normal samples are compared, and controlling for these biases will potentiate the discovery of molecular pathways and features otherwise hidden in the data.

To address this challenge we utilized two independent patient cohorts where the tissue composition of prostate cancer and normal samples has been thoroughly assessed by histopathology. Based on the gene expression analysis of stroma-enriched genes in these two cohorts, we used Gene Set Enrichment Analysis (GSEA)^25^ to assess the stroma content in five other patient cohorts where no histopathology is available. In total 1713 prostate cancer and 230 normal samples were assessed for their stroma content. To create datasets from all cohorts where the confounding effect of stroma tissue is accounted for, we used our recently published approach of balancing tissue composition^23^. When differential expression analysis is performed on these datasets, consistent downregulation of genes in the cholesterol synthesis pathway is highlighted as one of the most prominent features for primary prostate cancer.

## Results and Discussion

### Differentially expressed genes in seven publicly available prostate cancer cohorts controlled for stroma tissue confounding

We used seven publicly available cohorts of tissue samples from patients with prostate cancer (*Bertilsson, Chen, Taylor, TCGA, Prensner, Sboner* and *Erho*, referred to as the *seven-study-cohort*; N=1943 samples, 1713 prostate cancer and 230 normal, Table 1). Gene expression measurements in the various cohorts had been generated using different microarray platforms and RNA-sequencing. Of these seven cohorts, two cohorts (*Bertilsson* and *Chen*, referred to as the *histopathology cohorts,* Table 1) contained detailed histopathology on prostate cancer, stroma and benign epithelium in each sample. These two cohorts were used as a basis for stromal assessment in all seven cohorts. A flow-chart of the different steps in this assessment is provided in Figure 1, and a detailed description of each step is provided in the Methods section. Of the seven cohorts, five cohorts contained measurements of both prostate cancer and samples characterized as normal (*Bertilsson, Chen, Taylor, TCGA* and *Prensner*, referred to as the *five-study-cohort*; 1117 samples, 887 prostate cancer and 230 normal).

**Figure 1:**
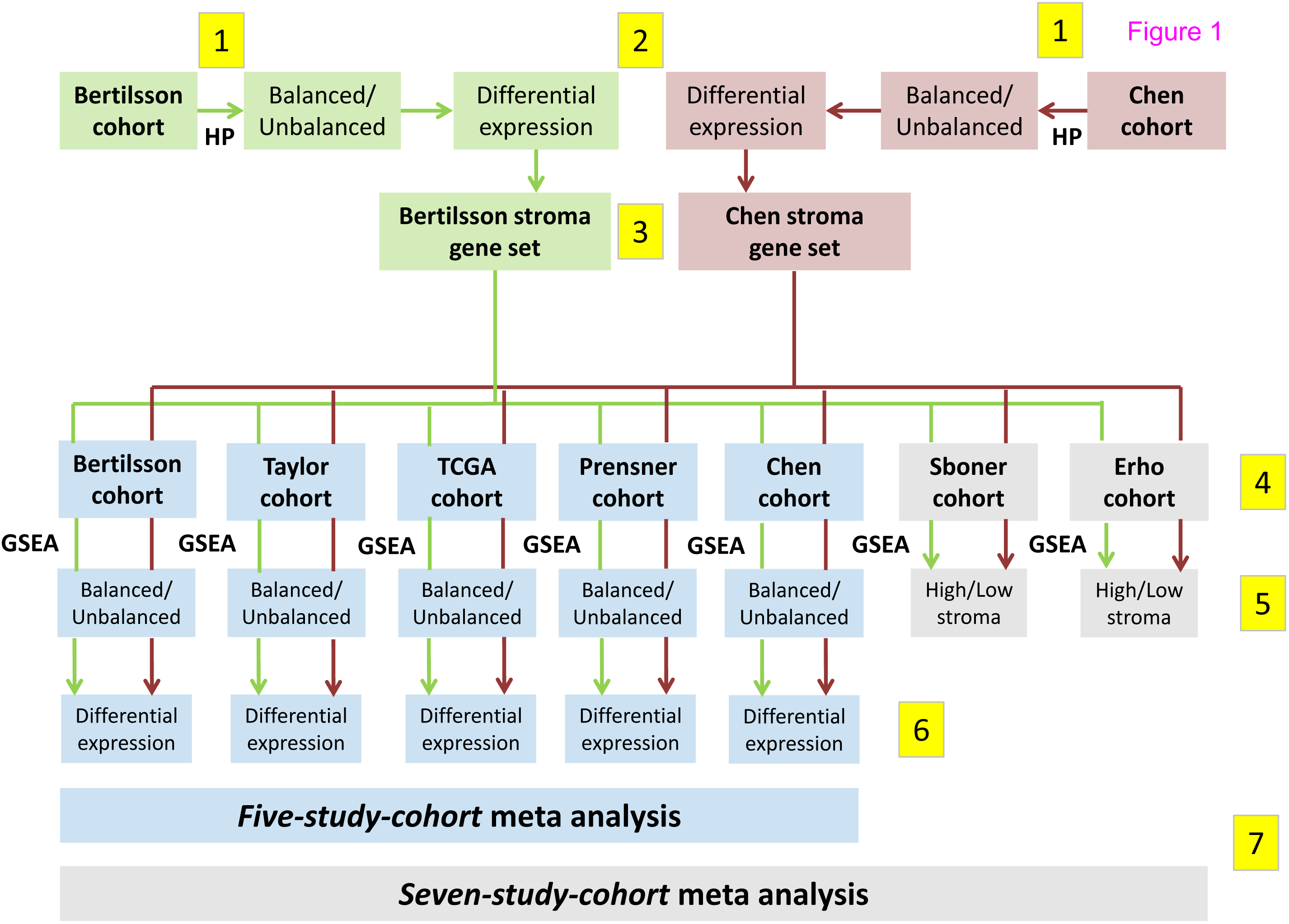
Flow chart illustrating the different computational steps for the analysis performed in this study. 1) Histopathology (HP) is used to create *balanced* and *unbalanced* datasets independently for the *Bertilsson* (marked green) and *Chen* (marked red) cohorts. 2) Differentially expressed genes for the HP-based *balanced* and *unbalanced* datasets are calculated for the *Bertilsson* and *Chen* cohorts. 3) Two stroma gene-sets are identified independently based on gene p-value relationships between the HP-based *balanced* and *unbalanced* datasets in the *Bertilsson* and *Chen* cohorts, respectively. 4) Gene Set Enrichment Analysis (GSEA) scores for all samples in all seven cohorts are calculated based on the two stroma gene-sets. These gene-sets are not combined, ensuring two independent GSEA stroma predictions for each sample in each cohort. 5) The GSEA scores are used to separate the five cohorts with both cancer and normal samples (including the cohorts from *Bertilsson* and *Chen*) into *balanced* and *unbalanced* datasets. The two remaining cohorts (*Sboner* and *Erho*) are only separated into groups with high and low stroma content. 6) Differentially expressed genes are calculated individually for the five cohorts with both cancer and normal samples. 7) *Balanced* and *unbalanced* datasets from the *five-study-cohort* are merged into one meta-analysis for differential expression. *Balanced* and *unbalanced* datasets from the *five-study-cohort*, as well as high and low stroma datasets from the *Sboner* and *Erho* cohorts are merged into one meta-analysis of the *seven-study-cohor*t.

**Table 1:** Data from the seven patient cohorts.

| Abbreviation | Source | Dataset reference | Article reference | Analysis Platform | Samples | PCa | Normal | Unique genes | HP on all samples |
| --- | --- | --- | --- | --- | --- | --- | --- | --- | --- |
| *Bertilsson* | Array Express | E-MTAB-1041 | 52 | Microarray, A-MEXP-2087 - Illumina Human HT-12 WG-DASL | 156 | 116 | 40 | 14 149 | Yes |
| *Chen* | GEO | GSE8218 | 21,53 | Microarray, Affymetrix Human Genome U133A Array | 136 | 65 | 71 | 12 497 | Yes |
| *Taylor* | GEO | GSE21034 | 54 | Microarray, Affymetrix Human Exon 1.0 ST Array | 160 | 131 | 29 | 18 294 | No |
| *TCGA* | TCGA | TCGA | 55 | RNA-Seq, Illumina HiSeq 2000/ Genome Analyzer IIX | 549 | 497 | 52 | 20 504 | No |
| *Prensner* | dbGaP | phs000443.v1.p1 | 56 | RNA-Seq, Illumina Genome Analyzer | 116 | 78 | 38 | 23 712 | No |
| *Sboner* | GEO | GSE16560 | 60 | Microarray, Human 6k Transcriptionally Informative Gene Panel for DASL | 281 | 281 | 0 | 6 102 | No |
| *Erho* | GEO | GSE46691 | 61 | Exon array, Affymetrix Human Exon 1.0 ST Array [probe set (exon) version] | 545 | 545 | 0 | 17 163 | No |

Footnote: HP=Histopathology; GEO= Gene Expression Omnibus; TCGA= The Cancer Genome Atlas; dbGap = The database of Genotypes and Phenotypes, PCa=Prostate Cancer

### Robust stroma assessment using Gene Set Enrichment Analysis (GSEA) with sets of stroma-enriched genes

A key concept in this study is to utilize GSEA^25^ assessments of stroma content in the tissue samples from the seven patient cohorts to create sub-datasets for differential expression analysis where the confounding effect of stroma tissue is accounted for^23^. To achieve this, we identified a robust and reliable stroma assessment protocol based on selecting genes which were up- or downregulated with respect to the content of stroma tissue in the two *histopathology cohorts* (Methods). The top ranked genes were collected into gene-sets used for GSEA-based assessment of stroma in samples from all seven cohorts. Stroma content in each sample was assessed independently by the two stroma gene-sets identified from the two *histopathology cohorts*. Although the two stroma gene-sets were generated from different patient cohorts using different microarray platforms, the genes identified showed on average 44∼ overlap for the top 1000 up‐ and downregulated stroma genes, compared to 6∼ for random genes (Figure 2a). A comparison with four previously published prostate stroma gene lists^21^, showed an average overlap of 62∼, compared to 8∼ for random genes (Figure 2b). Identified stroma genes were robust to two different methods for gene selection, with an average of 73∼ overlap (SFig2 in Supplementary File S1 online).

**Figure 2:**
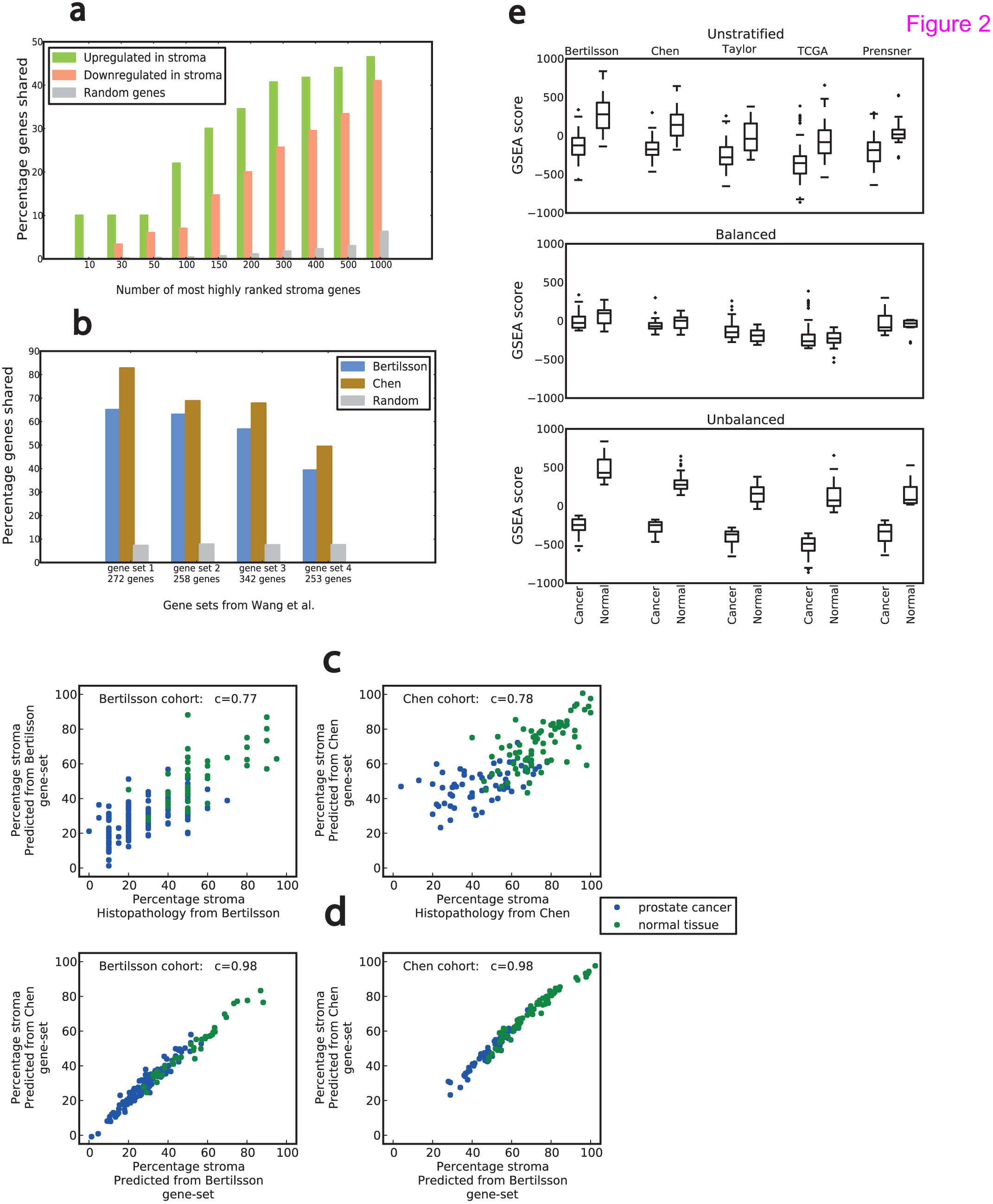
Robust assessment of stroma content in cohorts where no histopathology is available. a) Overlap between up- and downregulated stroma genes in the gene-sets from *Bertilsson* and *Chen* for various numbers of the N top-ranked stroma genes. The random numbers of shared genes are the average over 50 random gene selections for each N. b) Overlap of prostate stroma gene sets from Wang et al.^21^ with stroma gene-sets from *Bertilsson* and *Chen*. c) Pearson correlation (c) between predicted stroma percentage from GSEA and histopathological determined stroma percentage in the cohorts from *Bertilsson* and *Chen*. d) Pearson correlation (c) between stroma percentage predicted by gene-sets from *Bertilsson* and *Chen* in each of their respective cohorts (bottom). e) Bias towards higher GSEA stroma scores in normal compared to cancer samples present in all *unstratified* cohorts from the *five-study-cohort*. Dividing samples into *balanced* and *unbalanced* datasets minimizes and maximizes, respectively, the stroma bias between cancer and normal samples. A difference in the average overall GSEA score between the cohorts is also evident in the figure.

When varying the size of the stroma gene-sets, as well as the total number of genes used for the GSEA assessment, we observed an average deviation in predicted stroma content of only ∼1∼ (STab1 in Supplementary File S1 online). This shows that individual genes had minimal influence on the stroma assessments. In the two *histopathology cohorts*, the predicted stroma percentage from GSEA showed a mean deviation from histopathology between 10∼ and 11∼, (r=0.77 and r=0.78), respectively (Figure 2c). These measurements are in agreement with previously published comparisons between histopathology and gene based stroma predictions in prostate cancer ^21,26^. For the *histopathology cohorts*, the predicted percentages of stroma were highly correlated for the two gene-sets (r=0.98) (Figure 2c), and estimated GSEA scores in the additional five cohorts were also highly correlated (r between 0.95 and 0.99) (SFig3 in Supplementary File S1 online). GSEA scores for additional cohorts were also robust with respect to the size of the stroma gene-sets, with an average standard deviation for 0-100 normalized GSEA scores of ∼2 (STab1 Supplementary File S1online). Variations in GSEA scores were not dependent on the platform used for gene expression analysis. As further support for the validity of the stroma gene-sets, samples from several prostate cancer cell-lines included in the *Taylor* and *Prensner* cohorts, were consistently at the low end of stroma content when estimated by GSEA assessment. Overall, we conclude that the gene-sets give a stable, robust and reproducible representation of the stroma content in prostate cancer and normal tissue samples in each cohort. However, we observed a prominent baseline difference in the average GSEA scores between the different cohorts (Figure 2d). This means that an absolute prediction of stroma percentage for each sample which can be compared between cohorts cannot be made, but that relative stroma assessment between samples within the same cohort is feasible and robust.

### Balancing the stroma content in cohorts with missing histopathology

We used our previously published strategy^23^ (Methods) to create a stroma *balanced* and a stroma *unbalanced* datasets in each of the five cohorts with both cancer and normal samples. The balanced and unbalanced datasets are designed to have the same number of cancer and normal samples, making p-values from differential gene expression analysis directly comparable (STab2 in Supplementary File S1 online). In the *balanced* dataset, prostate cancer and normal samples have similar average stroma content, minimizing the tissue confounding commonly present in a conventional *unstratified* analysisDifferential analysis in the *balanced* dataset highlights changes between prostate cancer and normal tissue. In contrast, the *unbalanced* dataset is created to maximize the difference in stroma content between cancer and normal samples. In this setting, differentially expressed genes in prostate cancer compared to stroma will be highlighted. For a single gene, comparisons between the *balanced* and *unbalanced* datasets can reveal whether a significant differential expression truly results from changes between the normal tissue and cancer, or is due to variations in the average stroma content. Importantly, to create *balanced* and *unbalanced* datasets, no absolute estimation of stroma percentage in each sample is necessary. A relative stroma assessment is sufficient, ensuring that samples in the same cohort can be sorted according to their stroma content. This enables *balanced* and *unbalanced* analysis with our GSEA-based stroma assessments in cohorts where histopathology is not available.

We used the two stroma gene-sets identified independently from the *histopathology cohorts* (*Bertilsson* and *Chen)* to calculate GSEA scores for all 1943 samples (1713 cancer and 230 normal) from all the seven patient cohorts. In *the five-study cohort* containing 1117 samples (887 cancer and 230 normal), the calculated GSEA scores showed a systematic bias of increased average stroma content in normal samples (Figure 2d), which should support the separation of each cohort into *balanced* and *unbalanced* datasets. *Balanced* and *unbalanced* datasets were therefore created independently using the two available stroma gene-set, resulting in two independent *balanced* and *unbalanced* datasets for each cohort. This stratification equalized the average stroma content in the *balanced* datasets, and enhanced the difference in average stroma in the *unbalanced* datasets (Figure 2d). Differential expression analyses were performed independently for each dataset, and differentially expressed genes were ranked in each dataset according to their p-value. In addition, combined rank-based meta-analysis over the *five-study-cohorts* and *seven-study-cohorts* were performed (Methods). The *balanced* and *unbalanced* datasets from the two meta-cohorts contained 558/115 and 971/115 prostate cancer/normal samples each, respectively.

### Transcriptional downregulation of genes in the cholesterol synthesis pathway when adjusting for stroma tissue confounding

Consistent and highly significant downregulation of genes in the cholesterol synthesis pathway between cancer and normal samples was a prominent feature in the *balanced* analysis of gene expression (Figure 3a, Table 2, SFig4 in Supplementary File S1 online). In a meta-analysis of the *five-study-cohort*, 21 of the 25 genes assessed were downregulated.

**Figure 3:**
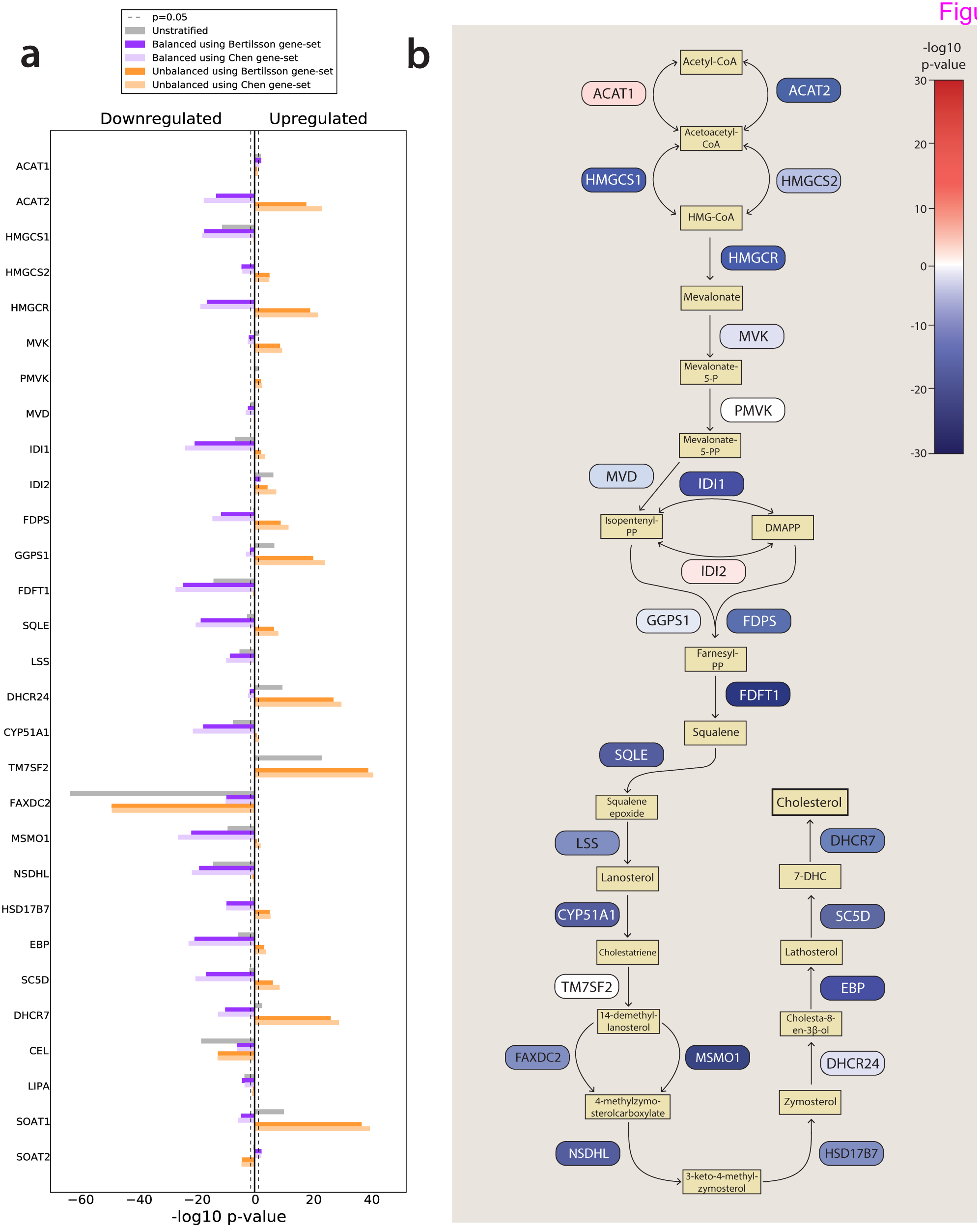
Genes in the cholesterol synthesis pathway are coherently downregulated in prostate cancer compared to normal epithelium. a) The figure shows –log10 p-values multiplied by 1 for upregulated genes, and −1 for downregulated genes. The results presented are for a rank-based meta-analysis of the *five-study-cohort*. All p-values presented are corrected for multiple testing using the total number of 25 964 unique gene identifiers from all cohorts. Results from individual cohorts as well as the *seven-study-cohort* can be found in SFig4 in Supplementary File S1 online. b) The schematic representation shows the cholesterol synthesis pathway with down- and upregulated genes color-coded in blue and red, respectively. The strength of the color corresponds to the degree of down- or upregulation.

**Table 2:** Overview of genes related to cholesterol synthesis assessed for differential expression.

| Gene symbol | Alternative symbol | Gene name | Gene function |
| --- | --- | --- | --- |
| ACAT1 |  | acetyl-CoA acetyltransferase 1 | Synthesis |
| ACAT2 |  | acetyl-CoA acetyltransferase 2 | Synthesis |
| HMGCS1 |  | 3-hydroxy-3-methylglutaryl-CoA synthase 1 | Synthesis |
| HMGCS2 |  | 3-hydroxy-3-methylglutaryl-CoA synthase 2 | Synthesis |
| HMGCR |  | 3-hydroxy-3-methylglutaryl-CoA reductase | Synthesis (rate limiting enzyme) |
| MVK |  | mevalonate kinase | Synthesis |
| PMVK |  | phosphomevalonate kinase | Synthesis |
| MVD |  | mevalonate decarboxylase | Synthesis |
| IDI1 |  | isopentenyl-diphosphate delta isomerase 1 | Synthesis |
| IDI2 |  | isopentenyl-diphosphate delta isomerase 2 | Synthesis |
| FDPS |  | farnesyl diphosphate synthase | Synthesis |
| GGPS1 |  | geranylgeranyl diphosphate synthase 1 | Synthesis |
| FDFT1 |  | farnesyl-diphosphate farnesyltransferase 1 | Synthesis |
| SQLE |  | squalene epoxidase | Synthesis |
| LSS |  | lanosterol synthase | Synthesis |
| DHCR24 |  | 24-dehydrocholesterol reductase | Synthesis |
| CYP51A1 |  | cytochrome P450 family 51 subfamily A polypeptide 1 | Synthesis |
| TM7SF2 |  | transmembrane 7 superfamily member 2 | Synthesis |
| FAXDC2 | C5orf4 | fatty acid hydroxylase domain containing 2 | Synthesis |
| MSMO1 | SC4MOL | methylsterol monooxygenase | Synthesis |
| NSDHL |  | NAD(P) dependent steroid dehydrogenase-like | Synthesis |
| HSD17B7 |  | hydroxysteroid (17-beta) dehydrogenase 7 | Synthesis |
| EBP |  | emopamil binding protein (sterol isomerase) | Synthesis |
| SC5D | SC5DL | sterol-C5-desaturase | Synthesis |
| DHCR7 |  | 7-dehydrocholesterol reductase | Synthesis (last step before cholesterol) |
| CEL |  | carboxyl ester lipase | Esterification |
| LIPA |  | lipase A, lysosomal acid, cholesterol esterase | Esterification |
| SOAT1 |  | sterol O-acetyltransferase 1 | Esterification |
| SOAT2 |  | sterol O-acetyltransferase 2 | Esterification |
| ABCA1 |  | ATP-binding cassette, sub-family A | Efflux |
| ABCG1 |  | ATP-binding cassette, sub-family G | Efflux |
| SLCO2B1 |  | solute carrier organic anion transporter family member 2B1 | Transport |
| SLCO1B3 |  | solute carrier organic anion transporter family member 1B3 | Transport |
| LDLR |  | low density lipoprotein receptor | Uptake |
| APOE |  | apolipoprotein E | Component for IDL, HDL and VLDL |
| SREBF1 |  | sterol binding element transcription factor 1 | Transcriptional activation |
| SREBF2 |  | sterol binding element transcription factor 2 | Transcriptional activation (main activator) |
| SCAP |  | SREBF chaperone | Transcriptional activation |
| MBTPS1 | S1P | membrane bound transcription factor peptidase site 1 | Transcriptional activation |
| MBTPS2 | S2P | membrane bound transcription factor peptidase site 2 | Transcriptional activation |
| INSIG1 |  | insulin induced gene 1 | HMGCR degradation |
| INSIG2 |  | insulin induced gene 2 | HMGCR degradation |
| AMFR | GP78 | autocrine motility factor receptor E3 ubiquitin protein ligase | HMGCR degradation |
| NR1H3 | LXRA | nuclear receptor subfamily 1<br>group H member 3 | Transcriptional repression |
| NR1H2 | LXRB | nuclear receptor subfamily 1<br>group H member 2 | Transcriptional repression |
| RXRA |  | retinoid X receptor alpha | Transcriptional repression |
| RXRB |  | retinoid X receptor beta | Transcriptional repression |
| MYLIP | IDOL | myosin regulatory light chain<br>interacting protein | Degradation of LDLR |

These included key genes of cholesterol synthesis such as *HMGCR* and *SQLE* (rate limiting enzymes), *FDFT1*, *LSS* (catalyzes first step), *DHCR7* (catalyzes last step) in addition to *NSDHL*, *MDMO1*, *EBP*, *IDI1*, *CYP51A1*, *HMGCS1* and *SC5D.* All these genes had p-values to the power of −10 or less. The same trend was observed in a meta-analysis over the seven-study-cohort (SFig4 in Supplementary File S1 online). In addition, four cohorts in the *five-study-cohort* individually showed highly significant downregulation of cholesterol genes, though the most highlighted genes varied somewhat between the cohorts (SFig4 in Supplementary File S1 online). In the *five-study-cohort*, 10 central cholesterol genes (*HMGCS1, HMGCR, IDI1, FDFT1, SQLE, CYP51A1, MSMO1, NSDHL, EBP* and *SC5D*) ranked among the top 150 most differentially expressed genes in the *balanced* dataset (average rank of 76) (Supplementary File S2 online). This is in contrast to the *unstratified* and *unbalanced* datasets, where the average ranks of the same ten genes were 9195 and 14860, respectively. The *unbalanced* analysis also shows that upregulation of cholesterol genes is mostly due to differences between cancer tissue and stroma (Figure3a). Cholesterol synthesis was a highly important gene ontology term in the *balanced* dataset, and a clustered set of related terms containing *steroid, sterol* and *cholesterol biosynthesis* were among the top three most significant gene ontologies when the 500 most significant genes from the *five-study-cohort* were analyzed by DAVID (Table 3). In summary, the *balanced* data prove a characteristic transcriptional downregulation of the cholesterol synthesis pathway in primary prostate cancer. All p-values presented in this and the following sections, as well as Figure 3 and Figure 4, are conservatively corrected for multiple testing using the total number of 25 964 unique gene identifiers from all cohorts.

**Figure 4:**
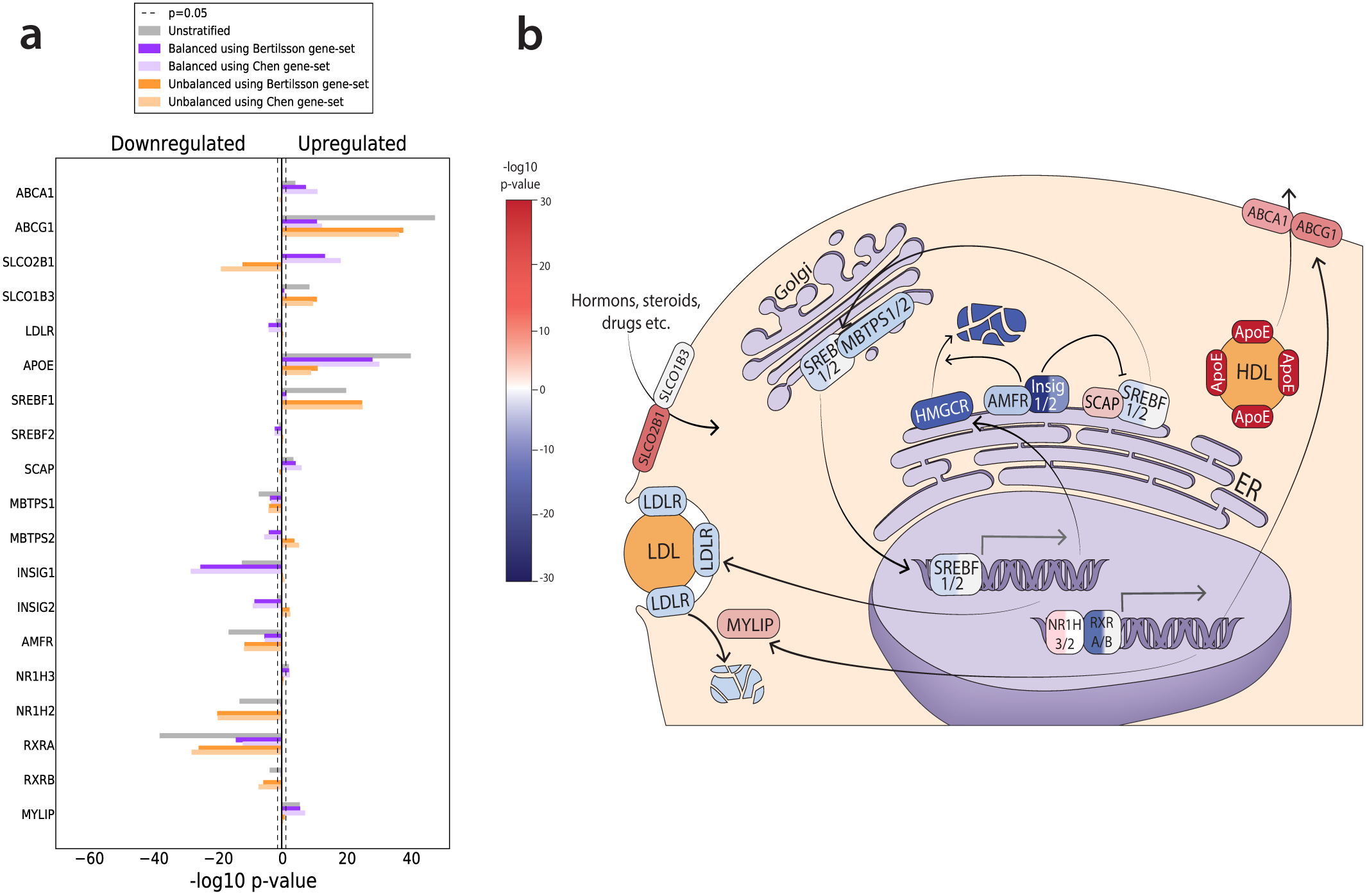
Differentially expressed genes from involved in cholesterol regulation, uptake, efflux and transport. Results from individual cohorts as well as the *seven-study-cohort* can be found in SFig5 in Supplementary File S1 online. a) The figure shows –log10 p-values multiplied by 1 for upregulated genes, and −1 for downregulated genes. All p-values presented are corrected for multiple testing using the total number of 25 964 unique gene identifiers from all cohorts. Results from individual cohorts as well as the *seven-study-cohort* can be found in SFig4 in Supplementary File S1 online. b) The schematic representation illustrates the cellular function of the selected genes, with down- and upregulated genes color-coded in blue and red, respectively. The strength of the color corresponds to the degree of down- or upregulation.

**Table 3:** Gene Ontology analysis identifies steroid, sterol and cholesterol biosynthesis among the most significantly altered pathways in prostate cancer.

| GO term | q-value using<br>*Bertilsson* gene-set | q-value using<br>*Chen* gene-set |
| --- | --- | --- |
| Cell Adhesion | 6.7e-8 | 6.6e-8 |
| Signal | 1.3e-7 | 1.6e-8 |
| Glycoprotein | 2.0e-8 | 1.1e-7 |
| **Steroid Biosynthesis** | **5.4e-6** | **4.3e-6** |
| **Cholesterol Biosynthesis** | **3.7e-4** | **1.7e-5** |

Footnote: The analysis was performed using the top 500 ranked genes from the *balanced* analysis in the *five-study-cohort* as input to DAVID. Only the top terms are listed. All terms are from the category “SP\_PIR\_KEYWORDS”. The top categories were the same when the top 1000 ranked genes were used. All terms related to steroid, sterol and cholesterol synthesis were part of the same functional cluster in DAVID.

### Expression of cholesterol pathway genes are confounded by stroma tissue

The pronounced discrepancies between the *balanced* and *unbalanced* datasets serve as an illustration of how cholesterol pathway genes are confounded by stroma tissue during differential analysis. Using *HMGCR* (the rate-limiting enzyme of cholesterol synthesis) as an example, this gene is strongly significant in both datasets. However, it is downregulated when cancer is compared to normal epithelium in the *balanced* dataset, and upregulated when cancer is compared to stroma in the *unbalanced* dataset. This typical pattern occurs when a gene highly expressed in the normal epithelium has an intermediate expression in cancer and is weakly expressed in stroma. Significant expression differences in these situations can only be revealed when the confounding effects of stroma is accounted for. Since this pattern is prevalent throughout the entire cholesterol pathway, we hypothesize that stroma confounding is the main reason that this pathway has not been identified in previous analysis of prostate cancer patient cohorts. The only cohort that did not highlight cholesterol synthesis was the cohort from *Chen*, which showed a consistent absence of significant cholesterol genes in the *balanced* dataset (SFig4 in Supplementary File S1 online). However, the cholesterol gene expression pattern from the *unbalanced* dataset in *Chen* was similar to the other cohorts.

### The selection of stroma genes does not cause bias on differential expression of cholesterol genes

It is important to establish that gene-sets representing stroma content do not impose unwanted biases with respect to the differential expression of cholesterol genes in additional cohorts. Here we present three arguments why this is unlikely for the cholesterol pathway genes in this study. 1) The stroma gene-sets were generated from two independent sources, but produced similar and stable results. 2) Cholesterol genes were either absent or ranked low in the stroma gene-sets. Nevertheless, all genes involved in the cholesterol pathway and regulation were excluded from any stroma gene-set during analysis to ensure unbiased sample stratification. Moreover, re-introduction of these cholesterol genes into the stroma gene-sets did not affect the stratification of samples into *balanced* and *unbalanced* datasets in any of the seven cohorts, showing that cholesterol-genes had no impact on the sample stratification. 3) The histopathology cohort from *Chen* was the only cohort that did not highlight cholesterol pathway genes as significant in the *balanced* dataset. Yet, all the *balanced* datasets in the other six cohorts still highlighted cholesterol genes as highly significant when the stroma gene-set derived from the *Chen* was used to balance the samples. Likewise, cholesterol pathway genes were not highlighted as significant in the *balanced* dataset from *Chen* when the stroma gene-set from *Bertilsson* was used to balance the samples in this cohort. This shows that both the *Chen* and the *Bertilsson* stroma gene-sets maintained the divergent *balanced* expression patterns for cholesterol genes when used in these two cohorts.

We also investigated two additional studies from the literature which could complement the findings in our study. One study emphasized cholesterol biosynthesis as a significant pathway in prostate tissue samples using gene ontology analysis ^27^. After analysis of the supplementary data material, genes in the cholesterol pathway showed negative cancer-to-normal fold-changes in that study (Supplementary File S3 online). The second study consisted of 50 samples (36 prostate cancer and 14 normal)^28^ collected using laser micro dissected tissue to avoid contamination from the stroma. Cancer-to-normal fold changes were negative for all key cholesterol genes in this study as well (Supplementary File S3 online). Thus the data in both these studies support the findings in our study.

### Decreased cholesterol synthesis may be beneficial for prostate cancer

Given the positive association between cholesterol and prostate cancer incidence, and the positive effect of statins on patient outcome, a consistent transcriptional downregulation of the cholesterol synthesis pathway in prostate cancer is a surprising observation. Although studies in prostate cancer cell-lines have demonstrated a role for cholesterol synthes is in tumor growth and aggressiveness, we have, after an extensive literature search, yet to see solid evidence for *in vivo* transcriptional upregulation of cholesterol synthesis in prostate cancer compared to normal tissue. Based on our results, we thus speculate how our observations may fit into the established mechanism of cholesterol metabolism for prostate cancer and for cells in general.

The regulation of cellular cholesterol levels is a highly complex and dynamic system, involving multiple feedback mechanisms (Figure 4b), where downregulation of cellular cholesterol synthesis is not necessarily contradictory to other observations. Cholesterol homeostasis in the cell is controlled by cholesterol synthesis, transport and storage, but the true *in vivo* balance between these sources has yet to be elucidated. The most established enzymes related to cholesterol homeostasis are *HMGCR* and *LDLR*^8^. *HMGCR* is the rate limiting enzyme for the cholesterol synthesis pathway in the cell, while *LDLR* controls the uptake of cholesterol from circulating Low Density Lipoprotein (LDL). In addition, the cell can store excess cellular cholesterol in prostasomes^4^ or by cholesteryl esterification in lipid droplets^9^. Increased availability of cholesterol from the environment may allow cells to shift their source of cholesterol from synthesis to uptake. Since cholesterol synthesis is energetically expensive^29^, this shift can be beneficial for the cancer cell to save energy, and a recent study in prostate cancer cell-lines showed that environmental cholesterol can supplement cellular cholesterol levels as a response to cholesterol synthesis inhibition^30^. Thus molecular precursors for cholesterol in the cell can be used in other pathways important for cancer growth. The shift may also prevent the anti-tumor activity of side products in the cholesterol pathway like oxysterols and isoprenoids, though the *in vivo* relevance for this mechanism is debated^30-33^. Additionally, the shift can provide an explanation why statins have a beneficial effect on prostate cancer patients. Statins mostly target cholesterol synthesis in the liver leading to reduced circulating levels of cholesterol^4^. This may limit the cholesterol available for cellular uptake, with activation of the cholesterol synthesis pathway and delayed cancer growth as a result. What contradicts this hypothesis is that not only *HMGCR* is downregulated in prostate cancer, but also *LDLR*. However, mechanisms alternative to *LDLR* for cholesterol and sterol uptake and efflux have been suggested, including changed activity of SLCO transporters^34^ (for example *SLCO2B1* is strongly upregulated in the *five-study-cohort*, Figure 4a) and modulation to cell-membrane structures like lipid rafts^3,7,35^. Recently, cholesteryl esters in lipid droplets in prostate cancer PC3 cells were shown to originate from uptake rather than synthesis^36^, supporting an increased attention to the role of cholesterol uptake in prostate cancer. Alternatively, statins may upregulate *HMGCR* in prostate cancer directly through feedback mechanisms^37^, again with a possible cancer-preventive effect. Finally, increased *HMGCR* protein levels have recently been shown to correlate with improved clinical outcome in breast^38^, colorectal^39^ and ovarian^40^ cancer. This may indicate that upregulation of the cholesterol pathway is a benign tumor characteristic, which is in line with the results presented here.

### Expression differences in regulatory genes suggest a possible compensation in cellular cholesterol synthesis by decreased HMGCR degradation

At the transcriptional level, *HMGCR* and *LDLR* mRNA are regulated, in particular by *SREBF2*, and partly by *SREBF1* transcription factors, which also regulates most of the enzymes in the cholesterol pathway^4,29,41^ (Figure 4b). SREBF is located on the membrane of the endoplasmic reticulum together with its cofactor *SCAP*. *SCAP* has a sterol-sensing domain, which activates SREBF-SCAP transport to the Golgi when sterol levels are low (Figure 4b). In the Golgi, SREBF-SCAP is enzymatically cleaved twice, which creates the nuclear active form of *SREBF1/2*. In the *balanced* dataset, *SREBF2* is downregulated while *SCAP* is upregulated (Figure 4). However, any potential increase in SREBF transport to the Golgi by *SCAP* is again counterbalanced by downregulation of both cleaving enzymes *MBTPS1* and *MBTPS2*. Thus the effect of *SCAP* upregulation on transcriptional activity is difficult to assess. *HMGCR* is also regulated at the translational level and at the level of degradation. We observe a strong downregulation of several key genes involved in *HMGCR* degradation ^41^, *INSIG1*, *INSIG2* and *AMFR* (Figure 4). Especially *INSIG1* is one of the most highly ranked genes differentially expressed in the *balanced dataset* (average rank 9). This suggests that some *HMGCR* activity can be maintained through downregulation of *INSIG1*, and that targeting *HMGCR* degradation can be an interesting option for modulating cholesterol levels in prostate cancer. Studies in model systems will be necessary to assess the combined effect of decreased transcription on one hand and decreased degradation on the other hand. The mechanisms of translational regulation of *HMGCR* are not well known, but may involve feedback regulation from side-products of the cholesterol pathway^31^.

Another pair of transcription factors implicated in negative regulation of cholesterol is the liver-X-receptors *NR1H3* and *NR1H2* (also called *LXRA* and *LXRB*) (Figure 4b), which dimerize with *RXRA* and *RXRB* to exert their regulatory activity^42^. *NR1H3* is upregulated in the *balanced* analysis (Figure 4), while its dimerization partner *RXRA* is strongly downregulated. We observe an upregulation of *NR1H3* targets, including the cholesterol efflux genes *ABCA1* and *ABCG1*, the *LDLR* suppressor *MYLIP* and a very strong upregulation of *APOE* (ranked highest among all genes in the *balanced* dataset). *APOE* can be an important constituent of High Density Lipoprotein (HDL) particles, where formation partly depends on the export by *ABCA1* and *ABCG1*. However, here our results are in disagreement with other *in vivo* reports, which associates low levels of *ABCA1* and low levels of circulating HDL with prostate cancer^43,44^. We finally emphasize that the discussion on how our data relate cholesterol metabolism and homeostasis are circumstantial, and that more detailed analysis involving proper model system is needed to elucidate these mechanisms further.

### Statin use and possible impact on downregulation of cholesterol synthesis pathway genes

A recent review reports that statins are ingested regularly by 25∼ of adults aged 45 years and older in the USA^17^. It is thus a possibility that statin use among the patients may have influenced the molecular makeup of the tumor at the time of surgery. We were not able to obtain sufficient data to conclude on this issue. Nevertheless, we here discuss the limited data and information we were able to find. Information on statin use prior to surgery were available only for the *Bertilsson* cohort, were a total of 26 samples (18 cancer and 8 normal) were affected. Re-analysis of the *Bertilsson* cohort did not change the pattern of consistent downregulation of cholesterol pathway genes (SFig6 in Supplementary File S1 online). There is one report on the *in vivo* effect of statin on HMGCR levels in breast cancer^37^. This report demonstrated that statins do not necessarily downregulate HMGCR, and that the effect of statin use was highly heterogeneous among patients. Currently we find it unlikely that statin use has a major impact on the highly significant and consistent results observed in our study, though we acknowledge that the information we have on this issue is too limited to conclude.

### Limitations to the histopathological tissue classification

In this study we have used a simplistic tissue classification which divide prostate cancer tissue into three tissue types; cancer, stroma and normal epithelium. However, this classification does not completely account for all tissue characteristics observed in prostate cancer, which can be heterogeneous with respect to all three tissue types. Cancer tissue from the prostate can be further classified into histological grades by Gleason score^45^. Gleason grading of samples was provided for six of the seven cohorts, and did not show any bias with respect to balanced and unbalanced dataset (STab2 in Supplementary File S1 online). We thus conclude that Gleason grade is not a confounding factor in our analysis. Several studies have shown that normal stroma can transform into reactive stroma when located adjacent to cancer tissue^46^. Thus the balanced analysis may also highlight genes resulting from differences between reactive and normal stroma. The strength of these differences will depend on the fraction of reactive stroma compared to normal stroma in the cancer samples. Histopathological differences between normal and reactive stroma were not assessed in the cohorts used in this study, and thus represents a limitation. Finally, normal epithelium from the prostate can display various precancerous aberrations with distinct molecular profiles^47^. We acknowledge that these are limitations of the current classification, and that further research and data generation in this field should focus on delineating additional molecular tissue profiles as well.

### Correlation between gene expression and protein levels

Finally, in this study, we have sometimes interpreted differences in gene expression of a gene as an indicator of protein level or activity, which is not necessarily related^48^. Nevertheless, transcriptional changes have been shown to be the most important mode of *HMGCR* and *LDLR* regulation, and the correlations between *HMGCR* and *LDLR* mRNA and protein level are comparable to mRNA and protein levels in general^49,50^.

## Conclusion

Analysis of differentially expressed genes between prostate cancer and normal samples in five patient cohorts, as well as meta-analysis over seven cohorts, consistently identified downregulation of nearly all genes in the cholesterol synthesis pathway in when the confounding effect of stroma tissue is minimized. This surprising observation will have important implications for our understanding of the complex relationship of prostate cancer and cholesterol metabolism.

## Methods

### Cholesterol pathway genes

Cholesterol genes were selected from KEGG^51^ pathway map for *Steroid Biosynthesis* and the *Mevalonate Pathway* in the pathway map for *Terpenoid Backbone Biosynthesis*. Twenty-five pathway genes were assessed for differential expression, which represent the complete pathways as mapped by KEGG. In addition, four genes from KEGG involved in cholesteryl ester formation and 19 genes from various literature sources involved in cholesterol regulation, uptake efflux and transport, were assessed. The complete list of genes and their main role in cholesterol homeostasis can be found in Table 2.

### Datasets, processing and quality assessment

**Data availability statement:** All gene expression data and associated metadata used in this study are publicly available in the database entries and references given in the data description below.

For expression analysis of genes in the cholesterol pathway we used gene expression measurements from prostate cancer and normal tissue samples from seven publicly available patient cohorts. An overview of data from the seven patient cohorts is given in Table 1. Cancer samples for all cohorts were from radical prostatectomy specimens, except for the *Sboner* cohort which was from a watchful waiting cohort. Normal samples were adjacent normal prostate tissue from prostatectomy specimens, except for four normal prostate samples in the *Chen* cohort which were autopsy samples from subjects without prostate cancer. Gene expression measurements from each patient cohort were downloaded and processed in the following way: Gene expression and metadata from *Bertilsson* were created by our group and processed as previously described^52^. Data are available at Array Express with accession E-MTAB-1041. The best probe for each gene was selected as the one with the highest average rank by p-value in differential expression analysis (average over *unstratified*, *balanced* and *unbalanced* comparison, see below or main text for explanation). Gene expression and metadata from *Chen*^21,53^ were downloaded from Gene Expression Omnibus (GEO) accession GSE8218. Probes were matched to gene names by the *hg133a.db* reference using *limma* in R. The best probe for each gene was selected as the one with the highest average rank by p-value in differential expression analysis. Gene expression and metadata from *Taylor*^54^ were downloaded from GEO accession GSE21034. Probes were matched to genes using the GPL10264 reference available at GEO. Probes with no matching gene were removed from further analysis. The best probe for each gene was selected as the one with the highest rank in a differential expression analysis between prostate cancer and normal samples. In the *Taylor* dataset, probes from the same gene generally had very similar ranks. Normalized and raw RNA-Seq read counts and gene names from *TCGA* where downloaded from The Cancer Genome Atlas [http://cancergenome.nih.gov],^55^. Normalized read counts were log2-adjusted before further analysis. For the *Prensner* cohort^56^, RNA-Seq raw reads in *fastq*-format were downloaded with approval from dbGap (project #5870) with accession phs000443.v1.p1. Raw reads were mapped to the hg19 transcriptome using *TopHat2*^57^, and *featureCounts*^58^ were used to assign the reads mapping to each gene. Normalization of gene counts were performed using the normalization formula from the *voom* program^59^. Gene expression and metadata from *Sboner*^60^ were downloaded from GEO with accession GSE16560. Probes were matched to gene names using the GPL5474 reference available at GEO. Only four genes in the *Sboner* cohort had more than one probe. For these genes, the probes with the highest overall expression value were selected as the best probe. Quantile normalized exon expression data and metadata from the *Erho* cohort ^61^ were downloaded from GEO with accession GSE46691. Exons identifiers were matched to gene names using the GPL5188 reference available at GEO. The total expression for each gene was calculated as the average expression over all exons for that gene. Differential expression of genes from *Bertilsson, Chen* and *Taylor* were identified using the *limma* package in R as described previously^52^, while *voom* on raw RNA-Seq read counts was used for differential expression of genes from *TCGA* and *Prensner*. In total, 1943 samples (1713 prostate cancer and 230 normal) with 25 964 unique gene identifiers were considered over all seven datasets (the *seven-study-cohort*). Five of the cohorts (*Bertilsson, Chen, Taylor, TCGA* and *Prensner,* referred to as the *five-study-cohort*) contained both prostate cancer and normal samples (in total 1117 samples, 887 cancer and 230 normal). The *seven-study-cohort* contained 4804 shared genes, and the *five-study-cohort* contained 9527 shared genes over their respective cohorts. Quality assessment of each cohort was performed by evaluating the Pearson correlations between genes in previously validated gene sets^25,62^ related to ERG-fusion, an established feature of primary prostate cancer ^63^ (SFig1 in Supplementary File S1 online). Samples from the *five-study-cohort* consistently displayed a higher average ERG-fusion gene correlation in prostate cancer samples compared to normal samples. Cancer samples in the *Erho* cohort showed a similar average correlation compared to cancer samples in the *five-study-cohort*, while the *Sboner* cohort showed weaker average correlation. Altogether six of the cohorts performed well for the quality assessment, while poorer quality was only indicated in the *Sboner* cohort.

### Stratification of a cohort into *balanced* and *unbalanced* datasets

For the stratification of samples into datasets of *balanced* and *unbalanced* stroma tissue composition we used a strategy recently developed in our research group^23^. The strategy can be applied to any cohort, as long as assessments of stroma content are available for both cancer and normal samples. We will here use the *Bertilsson* cohort to briefly describe the procedure (Figure 1). In the *Bertilsson* cohort, 116 prostate cancer and 40 normal samples were sorted independently according to their histopathologically determined percentages of stroma. From the sorted samples, two non-overlapping datasets were created by separating the cancer and normal samples into two equally sized groups. The *balanced* dataset pairs the 58 cancer samples with the highest percentages of stroma with the 20 normal samples with the lowest percentages of stroma, to create a dataset withequal average amounts of stroma in cancer and control samples. In contrast, the *unbalanced* pairs the 58 cancer samples with lowest percentage of stroma with the 20 normal samples with highest percentage of stroma, thus maximizing the difference of average stroma content between cancer and normal samples. The *balanced* dataset represents a comparison between prostate cancer and normal samples where the bias due to increased average stroma content in the normal samples has been minimized. Molecular differences in the *balanced* dataset are thus directly attributable to differences between cancer and normal tissue. The second dataset represent an *unbalanced* comparison where molecular differences mostly represent differences between prostate cancer and stroma tissue. Differentially expressed genes are then identified independently for the *balanced* and *unbalanced* datasets. The equal number of prostate cancer and normal samples in each dataset ensures a consistent statistical power, meaning that p-values are directly comparable for each gene between the *balanced* and *unbalanced* datasets. The number of samples used for balanced and unbalanced analysis in each cohort is provided in Supplementary STab2 in Supplementary File S1 online.

### Identification of gene-sets for assessment of stroma content in prostate tissue samples

Since the procedure for identification of gene-sets requires histopathology on both prostate cancer and normal samples in the same cohort, only the *Bertilsson* and *Chen* could be utilized for this purpose. Both these cohorts include detailed histopathological evaluation on the percentage tissue composition of prostate cancer, benign epithelium and stroma in both prostate cancer and normal samples. Stroma gene-sets were created independently from each of the two cohorts by the exact same procedure. The difference in average tissue composition between the *balanced* and *unbalanced* datasets (described in the previous section) facilitates the identification of genes specifically up- or downregulated in stroma compared to benign epithelium and cancer tissue by comparing p-values between the two datasets. Specifically, genes which display up or downregulation characteristic for stroma tissue will have lower p-values in the *unbalanced* compared to the *balanced* dataset. We thus used the following formula to rank all genes according to their suitability for creating stroma gene-sets:

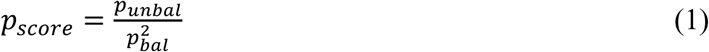

The squared term in the denominator was included to reflect that more pronounced differences in p-values are necessary for highly significant genes to be regarded as stroma genes. (Compare a gene with p-value 1e-5 in the *unbalanced* which is not significant in the *balanced* dataset, to a gene with p-value 1e-20 in the *unbalanced* and 1e-15 in the *balanced.* The former is more likely to be a valid stroma marker than the latter, even though the p-value ratio is the same). The stroma gene-sets included were based on the top 1000 ranked upregulated and top 1000 ranked downregulated genes. To avoid any bias from the cholesterol pathway genes, any genes from Table 2 were removed from all stroma gene-sets during analysis.

### Validation of stroma gene-sets in the *histopathology* cohorts

The independent stroma gene-sets created from the *Bertilsson* and *Chen* cohorts were validated by assessing the percentage of shared genes between the two gene-sets (Figure 2a). This percentage was compared to the percentage of shared genes expected by chance in 50 randomly generated gene sets of same size. The number of shared genes was also compared in gene sets created using a naïve approach of Pearson correlation to histopathological stroma content, and four previously published gene sets related to the content of stroma in prostate cancer tissue samples from Wang et al.^21^ (Figure 2b).

The assessment of the stroma content in any single sample from any cohort was performed using Gene Set Enrichment Analysis (GSEA)^54^. Two measures will influence the GSEA scores; the number of genes in the applied gene-set, and the total number of genes used for the calculation. We calculated 10 GSEA scores for each sample using varying numbers of the top scoring stroma genes (top 100, 150, 200, 250, 300, 350, 400, 450, 500 and 1000 genes), and normalized the scores in each of the 10 calculations to a 0-100 range over all samples to make them comparable. To enable comparisons between datasets, we only used genes shared by all datasets in each GSEA calculation. Two total gene selections were made, one containing 9527 genes shared by the *five-study-cohort*, and one with 4804 genes shared for the *seven-study-cohort*. Averaging over the 10 GSEA scores in each selection produced a total of four GSEA scores for each sample in *Bertilsson, Chen, Taylor, TCGA* and *Prensner* (using gene-sets from *Bertilsson* and *Chen* for the *five-study-cohort* and the *seven-study-cohort* respectively), and two GSEA scores for each sample in *Sboner* and *Erho* (*Bertilsson* and *Chen* gene-set for the *seven-study-cohort*). The main reason for the lower number of shared genes in the *seven-study-cohort*istherelativelyfewgenesmeasuredin*Sboner*(6100uniquegenes). For *Bertilsson* and *Chen,* GSEA scores for each sample were converted to predicted stroma percentages using a linear least squares fit, and compared to the stroma percentages obtained from histopathology. Predicted stroma percentages by the fit model based on the *Bertilsson* and *Chen* stroma gene-sets respectively in each of the two cohorts were also compared. Finally, GSEA score correlations when using the *Bertilsson* and the *Chen* stroma gene-sets were compared for all cohorts.

### Defining *balanced* and *unbalanced* data-sets in cohorts lacking histopathology

The GSEA scores representing the content of stroma in each sample were used to separate samples in each new patient cohort into *balanced* and *unbalanced* datasets as described above. This was done independently for each patient cohort. Differentially expressed genes for each cohort were calculated and corrected for multiple testing by Benjamini Hochberg false discovery rate (FDR) separately in each cohort, based on the total number of analyzed genes in each cohort. For the *Sboner* and *Erho* cohorts, only the cancer samples were separated into datasets with *high* and *low* stroma content, and no differential analysis was performed.

### Rank based meta-analysis for combined cohorts

To identify differentially expressed genes in a meta-analysis over the *five-study-cohort* the following procedure was used: 1) Each gene was sorted according to its expression value over all samples independently in each cohort, and rank-normalized to a score-value between 0 and 100, where 0 is the rank based expression value for the sample with the lowest expression value of the gene, and 100 is rank-based expression value the sample with the highest expression. 2) Rank-normalized values were mean centered independently in each cohort, where the mean centering was weighted by the relative number of prostate cancer and normal samples in the cohort. This was to avoid mean-value biases due to the huge relative difference between cancer and normal samples in each cohort. 3) Samples were separated into one *balanced* and one *unbalanced* meta-dataset using their previous assignment to *balanced* and *unbalanced* datasets in each individual cohort. Differential analysis based on two classifications were performed, one based on the stroma gene-set from *Bertilsson* and one on the stroma gene-set from *Chen*. 4) Differentially expressed genes for the *unstratified*, *balanced* and *unbalanced* datasets were calculated for weighted mean centered rank-normalized values between prostate cancer and normal samples combined for all five cohorts using the Mann-Whitney-Wilcoxon test^64^ for rank-based differential expression. P-values of differentially expressed genes were corrected for multiple testing using the Benjamini-Hochberg FDR for the total number of genes analyzed (25 964 unique gene identifiers). If a gene was not present in all datasets, only the datasets that contained that gene were used for differential expression. The *seven-study cohort* was analyzed in the same way, but with mean centering rather than weighted mean centering used to adjust gene-ranks between cohorts. This was due to the lack of normal samples in the *Sboner* and *Erho* cohorts.

### Gene ontology analysis

The top 500 and top 1000 differentially expressed genes from the rank-based differential expression analysis based on the both the *Bertilsson* and *Chen* gene-sets (four lists of genes in total) were subjected independently to DAVID ^65^ for gene ontology analysis.

### Ethical statement

The use of human tissue material and clinical data from the *Bertilsson* cohort was approved by the Regional Committee for Medical and Health Research Ethics (REC) for Central Norway, approval no 4-2007-1890. All experiments were performed in accordance with relevant guide lines and regulations. Informed consent was obtainedfromallparticipants.

Other ethical aspects regarding the specific samples used in this study have been described in previous publications^52,66^. RNA-Seq data from the *Prensner* cohort was approved through dbGap (project #5870), and data were downloaded and stored according to the provided security requirements. All other data were downloaded from freely available and publicly open resources.

## Acknowledgements

This works supported by the Liaison Committee between the Central Norway Regional Health Authority (RHA) and the Norwegian University of Science and Technology (NTNU) to [MBR]; the Norwegian Cancer Society [100792-2013] to [TFB], PhD position from Strategic funding ISB, Norwegian University of Science and Technology (NTNU) to [MKA], PhD position from Enabling Technologies, Norwegian University of Science and Technology (NTNU) to [KR]. The technique for fresh frozen tissue biobanking and cylinder extraction for reference E-MTAB-1041 was developed by Biobank1, St.Olavs Hospital, Trondheim, Norway. Funding support for MPC_Transcriptome sequencing to identify non-coding RNAs in prostate cancer was provided through the NIH Prostate SPORE P50CA69568, R01 R01CA132874, the Early Detection Research Network (U01 CA111275), the Department of Defense grant W81XWH-11-1-0331 and the National Center for Functional Genomics (W81XWH-11-1-0520). The results shown here are in part based upon data generated by the TCGA Research Network: http://cancergenome.nih.gov.

## Author Contributions

MBR, FD and MBT conceived the idea and developed the concept. MBR, MBT, HB and MKA performed data curation. MBR KR and FD performed analysis. MBR and FD developed the method. MBR, MBT, FD and TFB acquired funding and performed supervision. MBR, MBT, FD, HB, and TFB prepared the original draft. MKA and MBR prepared figures. All authors wrote and reviewed the manuscript.

